# Ant queens cannibalise infected brood to contain disease spread and recycle nutrients

**DOI:** 10.1101/2024.03.13.584778

**Authors:** Flynn Bizzell, Christopher D. Pull

## Abstract

Filial cannibalism, where parents eat their own offspring, is a taxonomically widespread behaviour with a multitude of potential adaptive explanations. Of these, the impact of parasites on the expression of filial cannibalism is particularly poorly understood. On one hand, cannibalising young with low survival probability may enable parents to reinvest valuable resources into future reproduction; on the other, cannibalising offspring harbouring parasites that can potentially also infect the parents may select against this behaviour. Although disease-induced cannibalism of eggs has been reported in fish, the benefits of consuming infected brood to contain infections – as an explanation for the evolution of filial cannibalism – remains largely unexplored. Here, we demonstrate that solitarily founding ant queens cannibalise all sick larvae in their nests before they become contagious, showing clearly that filial cannibalism both (i) contains an otherwise lethal infection without any long-term consequences on queen survival and (ii) enables the reinvestment of recouped energy into additional egg production.

## Main text

Filial cannibalism, where parents eat their own offspring, is a taxonomically widespread behaviour with a multitude of potential adaptive explanations [1]. Of these, the impact of parasites on the expression of filial cannibalism is particularly poorly understood. On one hand, cannibalising young with low survival probability may enable parents to reinvest valuable resources into future reproduction [1]; on the other, cannibalising offspring harbouring parasites that can potentially also infect the parents may select against this behaviour. Although disease-induced cannibalism of eggs has been reported in fish [2], the benefits of consuming infected brood to contain infections – as an explanation for the evolution of filial cannibalism – remains largely unexplored. Here, we demonstrate that solitarily founding ant queens cannibalise all sick larvae in their nests before they become contagious, showing clearly that filial cannibalism both (i) contains an otherwise lethal infection without any long-term consequences on queen survival and (ii) enables the reinvestment of recouped energy into additional egg production.

Ant queens found colonies alone and most fail due to predation, starvation, or infection of the queen or her brood [3]. Although the collective disease defences performed by workers in mature social insect colonies – termed social immunity – are well characterised [4], we still know surprisingly little about the behavioural immunity of their founding individuals [5]. Here, we report that founding queens of the black garden ant (*Lasius niger*) have evolved a unique yet effective response towards infected young. Specifically, we exposed five larvae per queen to infectious conidiospores of the fungal pathogen *Metarhizium* (strain isolated from a naturally infected *L. niger* queen). Larvae were left in isolation for 24 hours to develop lethal, but not yet transmissible, infections. As a control, larvae were exposed to a sham treatment and similarly isolated. After 24 hours, all larvae were returned to the queens and their response filmed. In total, we found that 92% of infected larvae were fully cannibalised by the queen leaving no remains – whereas just 6% of control larvae were consumed (Figure 1A and Movie S1; Likelihood ratio test (LR)-χ2 = 89.23, df = 1, *p* < 0.0001; n = 10 queens in both treatments).

**Figure 1.**
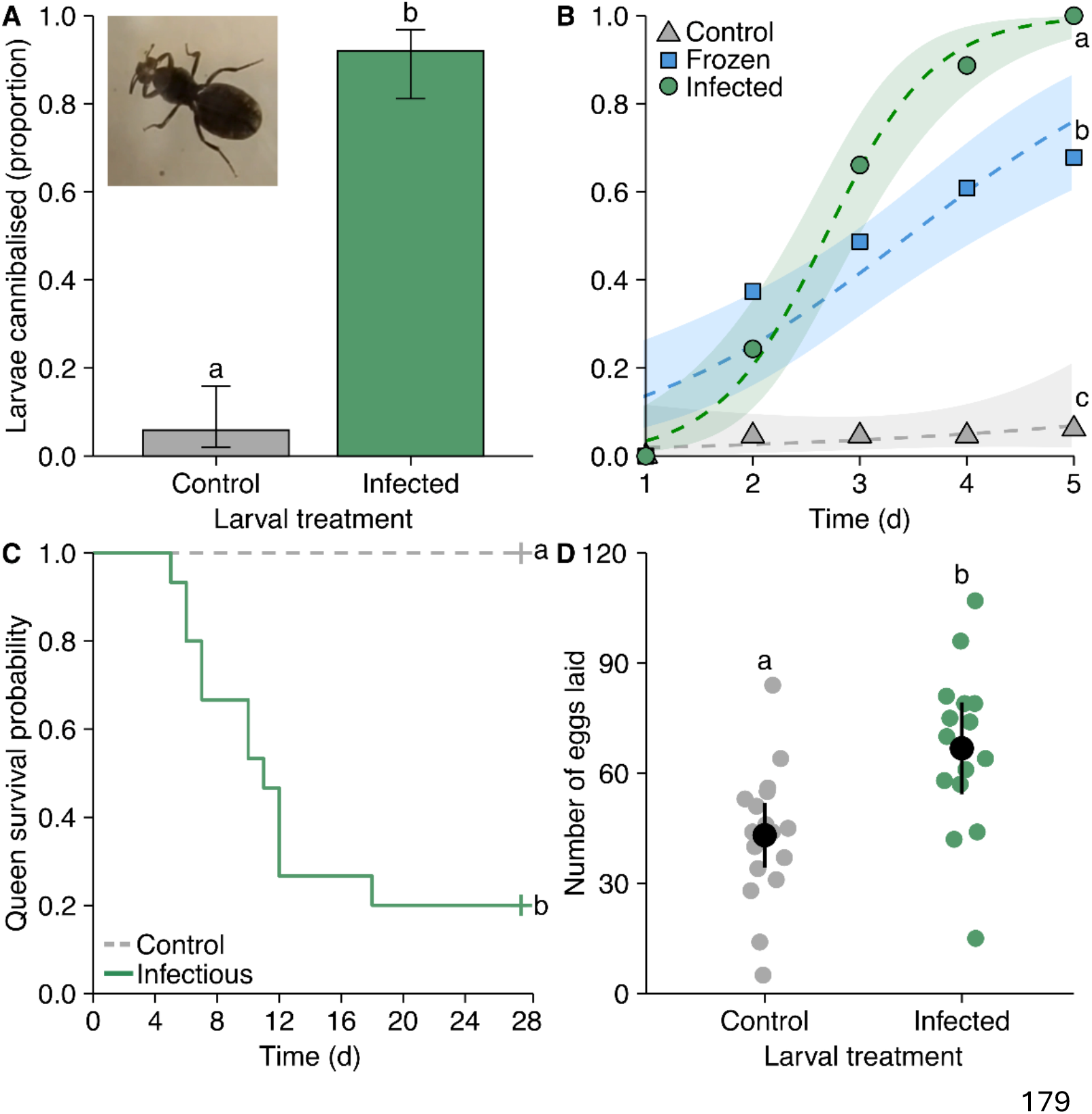
Cannibalism of infected brood by ant queens. (A) Ant queens cannibalise more larvae infected with the fungal pathogen *Metarhizium* than sham treated controls. Inset shows ventral view of queen eating infected larva; bars show proportion ± 95% confidence intervals (CI). (B) Infected larvae are cannibalised at faster rates compared to control or freeze-killed larvae; lines represent logistic regression curves ± 95% CI. (C) Queen survival probability is drastically reduced when they fail to cannibalise infected larvae; crosshairs denote surviving queens. (D) Queens that cannibalise infected larvae subsequently lay more eggs than control queens that do not perform cannibalism; error bars show mean ± 95% CI and each point represents one replicate. (A-D) Letters denote significantly different groups (*P* < 0.05).

To test if cannibalism is a response to infection or simply the death of the larvae, we presented queens with either control, infected, or freeze-killed, non-infected larvae, finding that infected larvae were cannibalised at substantially higher rates (Fig 1B; interaction between treatment and time: LR-χ^2^ = 65.09, df = 2, *p* < 0.0001; *p* ≤ 0.0003 for all post-hoc comparisons; n = 26 control and 23 infected/freeze-killed). Social insect queens, like their workers, are thus able to detect early-stage brood infections before they become infectious [4]. Despite the potential risk of infection, we surprisingly detected no long-term consequences of cannibalism on queen mortality, with the majority surviving the solitary period and producing workers (Fig S1A). Moreover, final colony size and composition did not differ between treatments (Fig S1B). Ant queens may minimise their risk of contracting infections via cannibalism by swallowing their acidic, antimicrobial venom, to create a gut environment that neutralises pathogens, as reported in workers [6]. Indeed, we anecdotally observed queens grooming the abdominal venom gland opening.

Next, to understand why queens cannibalise their infected young, we simulated a failure to detect and cannibalise brood by presenting queens with infectious larvae (sporulating cadavers covered in thousands of conidia) or control larvae. Queens intensively groomed and sprayed infectious cadavers with antimicrobial venom (Movie S2) but despite these apparent attempts to contain the infection, only 20% of queens survived in the infectious cadaver group (Figure 1C; infectious vs control: LR-χ2 = 20.3, df = 1, *p* < 0.0001; n = 15 queens in both treatments). Even when the queen did survive, all her healthy brood was lost to secondary infections.

“Energy benefit” explanations for the evolution of filial cannibalism predict parents should consume their brood under unfavourable conditions, so recouped resources can be redirected into future reproduction [1]. However, it is unknown whether infections limit the nutrient recovery process. We found that queens who consumed their infected larvae laid an additional 55% more eggs than non-cannibalising control queens (Figure 1D; *t* = -3.30, df = 26.45, *p* = 0.003; n = 15 infected and 18 controls). Reinvesting recouped nutrients back into egg production may be especially critical for founding queens: they do not forage, surviving solely on the breakdown of bodily reserves, so are severely resource limited [7]. Producing enough brood is essential as intraspecific competition with other young, neighbouring nests is a major driver of incipient colony failure, and larger colonies invariably win [7].

Combined, our results reveal that filial cannibalism by ant queens is an induced response to infection that prevents disease causing colony founding failure. Given how readily ant queens perform “hygienic cannibalism”, it is surprising worker ants in mature colonies do not express this behaviour. Two potential explanations for this difference are: firstly, unlike workers [8], queens cannot readily dispose of infected brood by discarding it outside of the nest, as they seal themselves into their claustral cell. Secondly, queens naturally consume a high protein diet to fuel egg production, whereas worker lifespan is shortened by consuming too much protein [9]. However, cannibalism of both healthy, infected, and dead nestmates by termite workers is commonly observed [10]. Like founding ant queens, many termites live in “closed” nests, precluding extranidal corpse disposal, and their high cellulose diet makes other nestmates a rare source of protein. Consequently, founding ant queens and termite workers may have convergently evolved filial and nestmate cannibalism as distinct strategies to solve the same problem of disease containment in a confined space, whilst simultaneously ensuring valuable nutrients are not wasted.

In conclusion, our study reveals that preventing the spread of infections among brood, coupled with the beneficial recovery of resources from sick offspring, can drive the evolution of filial cannibalism; a previously predicted but hitherto untested hypothesis [2]. Additionally, we find social insect queens are remarkably effective at detecting and eliminating brood infections before they become transmissible, providing further support for the theory that worker social immunity evolved from maternal care [4].

## Acknowledgements

This research was supported by the Oxford University Press John Fell Fund (award number 0009653).

## Author contributions

F.B. and C.D.P. designed the study and wrote the manuscript. F.B. performed experiments and data analysis with input from C.D.P.

## Supplementary Information

Supplementary methodology, figures (Fig. S1), movies (Movie S1-2), and additional references.

## Deposited Data

All data supporting this work has been uploaded to FigShare (link for review: https://figshare.com/s/1b9d1200ac491a9efaaf).

