## Supplementary Information for "Ant queens cannibalise infected brood to contain disease spread and recycle nutrients"

### Supplemental information

#### Supplemental figures

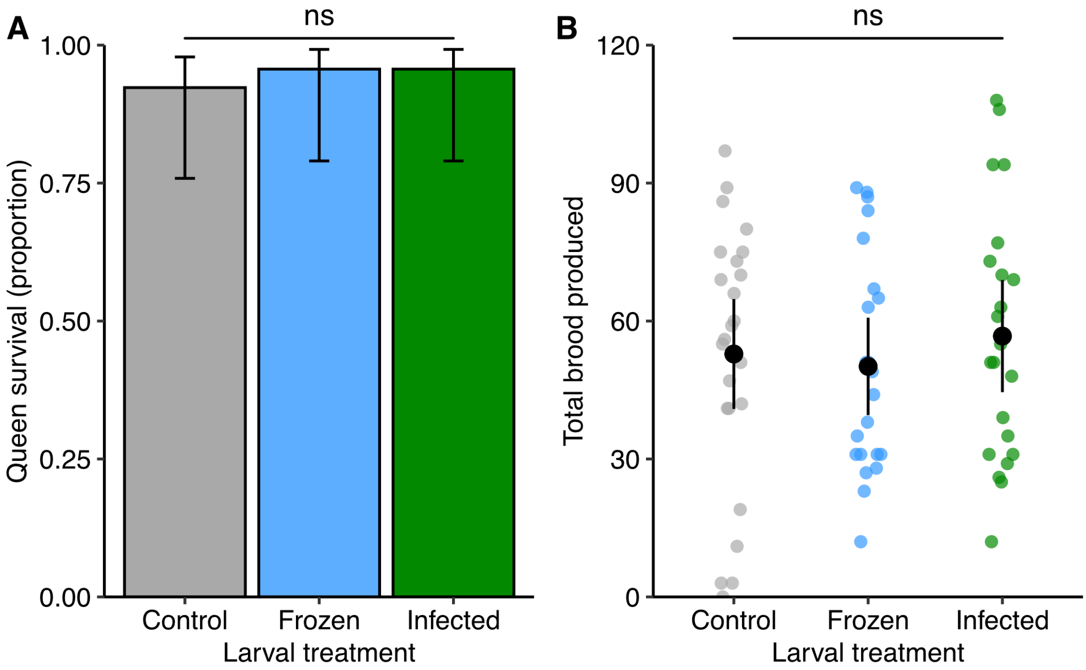


**Figure S1. Long term queen survival and brood production.** (A) Ant queens that cannibalised infected larvae have similar survival rates to control queens (LR-$\chi^2$ = 0.34, df = 2, $p$ = 0.84); bars represent proportion of queens surviving ± 95% confidence intervals (CI). (B) Ant queens that cannibalise infected larvae have an overall comparable number and composition of eggs, larvae, pupae and workers to control queens at the end of the colony founding period (perMANOVA $F_{2,65}$ = 0.35, $p$ = 0.95); error bars show mean total number of eggs, larvae, pupae and workers combined ± 95% CI and each point represent one replicate. (A-B) ns = no significant difference between treatments.

#### Supplemental Movies

**Movie S1.** Queen filmed from beneath cannibalising infected larvae at approx. x10 speed.

**Movie S2.** Queen grooming and spraying with acidic venom a sporulating larva.

**Supplemental experimental procedures**

*Queen collection and maintenance*

Hundreds of mated queens (identifiable by the loss of wings) were collected during a large nuptial flight in Oxfordshire, UK in July 2023. Queens were temporarily housed together in plastic boxes containing damp tissue paper. On return to the lab, each queen was weighed using a microbalance and placed alone into a petri dish with a damp plaster substrate, imitating a claustral cell. Queens were kept at 21°C in a temperature-controlled room on a 12-hour day/night light cycle and dishes were moistened weekly to maintain sufficient humidity levels. No food was provided to the queens as foundresses survive solely on the histolysis of the redundant wing muscles and fat reserves. Queens were chosen for each experiment/treatment group haphazardly.

*Fungal pathogen*

*Metarhizium brunneum* is a generalist, opportunistic entomopathogenic fungus known to infect many insect species [1] including ants and founding queens of our study species [2,3]. We used a strain that was isolated from a naturally infected founding *L. niger* queen. *Metarhizium* spp. are abundant in the environment as pathogenic asexual conidiospores (dispersal stage; ~ 5000 infectious conidiospores/g of soil [S4]); when conidiospores come into contact with a host's cuticle, they germinate and penetrate into the hemocoel to establish an internal infection. Infections remain non-transmissible until they eventually kill the host and sporulate out of the corpse, producing thousands of secondary conidiospores that can infect new insects [1]. The fungus was grown from long-term cryogenic stocks on sabaroud dextrose agar plates before each experiment. Conidiospores from fully sporulating plates were harvested and suspended in autoclaved 0.05% Triton X-100 and washed three times by centrifuging the conidiospores, pouring away the supernatant, and re-suspending them again in sterile 0.05% Triton-X. The viability of the conidiospores was confirmed by plating out 100 µl of the conidiospore suspension onto sabouraud dextrose agar plates and checking the proportion germinating after 18 h (always >90%).

*Larval fungal exposure and infection timeline*

To generate infected larvae for all experiments, we removed five similarly sized larvae per queen, placed them onto a strip of ethanol-wiped parafilm, and then rolled all five larvae simultaneously in 1µl of either the fungal conidiospore suspension or autoclaved 0.05% Triton-X as a sham control. We used a conidiospore concentration of $10^8$ conidia ml$^{-1}$ throughout the study. Larvae from the same queen were grouped and left in isolation for 24 h in new plaster-based petri dishes to allow infections to develop, before use in experiments. In a preliminary study, we removed a pair of larvae from ten different queens and exposed one to the conidia suspension (n = 10) and the other the control solution (n = 10), following the same procedure as above. We confirmed that all larvae exposed to conidia and left in isolation developed lethal infections: major hyphal growth was observed two days after exposure and all larvae had sporulated by day five. In contrast, all control larvae were still alive on day five, determined by gently prodding each larva with a toothpick for 10 s and monitoring for "wiggling" movement.

*Queen response to infected larvae*

To determine how queens respond to infected larvae, five larvae per queen were removed and exposed to the pathogen or control as above. Following the 24 h isolation period, the queen was placed into the petri dish with her now infected or control larvae (n = 10 queens in both). Queens were added to the larval dish rather than the other way around as treated and nontreated brood are indistinguishable. Petri dishes were gently flipped upside down so that the brood and queen were repositioned onto the lid, with the solid plaster base above them. The upside-down dish was then suspended on a clear, acrylic platform above a camera; this arrangement allowed us to observe the queen's interactions with her brood from underneath without her body obscuring the view. We recorded each petri dish for 24 h using a Raspberry Pi 4 running "PiSpy" software for animal behavioural monitoring [4], connected to a High-Quality Raspberry Pi camera with a 3MP 8-50mm 1/2.5" F1.4 C mount lens. Videos were then observed to establish queens were cannibalising the infected larvae.

*Queen cannibalism of control, dead, and infected larvae, and long-term colony founding success*

To test if larval cannibalism by queens is a response to infection or simply larval death, we generated infected and control larvae as above; this time, however, we added a third treatment group comprising dead larvae that had been freeze-killed for several hours at –70˚C and then defrosted. Queens were added to the petri dish as before (n = 26 control, 23 infected, and 23 frozen). We then recorded the number of larvae cannibalised over a five-day period. Following this five-day period, queens were returned to their original petri dishes and brood (no noticeable mortality was observed among original brood in the queens' absence). We then monitored dishes weekly for queen mortality until most queens had produced their first workers. A colony count was made at the end of the experiment (~ 5 months post-cannibalism), recording the final numbers of eggs, larvae, pupae, and workers the queens had produced.

*Survival of queens and healthy brood confronted with sporulating larvae*

To better understand the benefits of cannibalism, we simulated a scenario in which the queens "failed" to cannibalise infected larvae and monitored subsequent secondary disease transmission. Infected larvae were produced as above, but this time left in isolation for six days until all larvae had died and sporulated, and so become infectious. Control larvae were simply exposed to Triton-X but only kept in isolation for one hour. Both groups of larvae were then returned to the original petri dishes (n = 15 in both treatments), containing the queen and the rest of her healthy brood. The survival of queens and healthy brood was then monitored daily for 28 days (when queen mortality plateaued).

*Queen egg laying following cannibalism*

We tested the hypothesis that cannibalism allows queens to recycle nutrients by monitoring the number of eggs queens laid post-cannibalism. Using the same set up as before, queens were placed into new petri dishes containing five infected larvae to cannibalise (n = 15). As a control, queens were placed into new petri dishes but given no larvae (n = 18). The number of eggs laid by both cannibal and control queens was then counted using a microscope for 14 days; as before, all infected larvae were cannibalised by the queens within a few days.

*Data Analysis*

All statistics were performed using R v4.3.1 [5] and data has been published on FigShare [6]. We analysed the proportion of larvae cannibalised by each queen (Fig. 1A) using a logistic regression (general linear model with binomial error distribution and log-link function). We included the number of larvae cannibalised and the total number of larvae initially given to queens as the response, using the R *cbind* function to create a matrix of 'successes' and 'failures' directly within the model; larval treatment (control/infected) was included as the only fixed effect. We compared the full model to a null (intercept-only) model to assess the effect of the predictor, using likelihood ratio tests [7]. The proportion of larvae cannibalised over time across treatments (Fig. 1B) was analysed using a mixed effects logistic regression (generalised linear mixed effects model with binomial error structure and log-link function). Again, we used the *cbind* function to include the proportion of larvae cannibalised as a response matrix, with larval treatment (control/frozen/infected), day (*z*-transformed), and their interaction as fixed effects. To control for the repeated observation of the same replicate over time, we incorporated random intercepts for each replicate that interacted with random slopes for day of experiment, enabling us to explicitly model individual differences over time. Again, we compared the full model to a null (intercept-only) model to first assess goodness-of-fit ($p < 0.0001$), followed by a reduced model without the interaction, using likelihood ratio tests. We performed post hoc analyses using the *emtrends* function [8] to estimate and compare the slopes of the relationship between each level of treatment and day, correcting the resulting $p$ values using the Benjamini-Hochberg procedure for multiple testing [9]. We also used a logistic regression (general linear model with binomial error distribution and log-link function) to analyse the long-term survival of queens (Fig. S1A), including queen mortality (dead/alive) as the response and treatment as the only predictor; we again compared the full model to a null (intercept-only) model using likelihood ratio tests. For all logistic regressions, the appropriate assumptions/model diagnostics were checked, including the presence of model instability, influential observations, overdispersion, and multicollinearity (when multiple predictors present). We compared the final size and composition of colonies between treatments (Fig. S1B) using a non-parametric perMANOVA analysis of Mahalanobis dissimilarities, which is robust to multivariate covariance and makes no assumptions about the normality/variance of the data. This approach compares the overall multivariate structure/composition of the data among treatment groups; we included the number of eggs, larvae, pupae, and workers as a multivariate response matrix and treatment (control/frozen/infected) as the only predictor. We used a log-rank test to compare the survival curves of queens presented with either infectious or control larvae (Fig. 1C); queen survival was included as a survival object and larval treatment (control/sporulating) as the only fixed effect. Finally, we analysed the number of eggs laid by queens (Fig. 1D) using a two-sample *t*-test, with the number of eggs laid as the response and larval treatment (control/infected) as the fixed effect, having ensured the data met the assumptions of normality and homogeneity of variances (via visual inspection of histogram and a Levene test). Throughout, we used a combination of the packages: *binom* [10], *car* [11], *DHARMa* [12], *lme4* [13], *patchwork* [14], *performance* [15], *survminer* [16], *survival* [17], *tidyverse* [18], *vegan* [19].
